# yacrd and fpa: upstream tools for long-read genome assembly

**DOI:** 10.1101/674036

**Authors:** Pierre Marijon, Rayan Chikhi, Jean-Stéphane Varré

## Abstract

**Motivation:** Genome assembly is increasingly performed on long, uncorrected reads. Assembly quality may be degraded due to unfiltered chimeric reads; also, the storage of all read overlaps can take up to terabytes of disk space.

**Results:** We introduce two tools, `yacrd` and `fpa`, preform respectively chimera removal, read scrubbing, and filter out spurious overlaps. We show that `yacrd` results in higher-quality assemblies and is one hundred times faster than the best available alternative.

**Availability:** https://github.com/natir/yacrd and https://github.com/natir/fpa

**Supplementary information:** Supplementary data are available online.

## 1 INTRODUCTION

Third-generation DNA sequencing (PacBio, Oxford Nanopore) is increasingly becoming a go-to technology for the construction of reference genomes (*de novo* assembly). New bioinformatics methods for this type of data are rapidly emerging.

Some long-read assemblers perform error-correction on reads prior to assembly. Correction helps reduce the high error rate of third-generation reads and make assembly tractable, but is also a time and memory-consuming step. Recent assemblers (e.g. Li (2016); Ruan and Li (2019) among others) have found ways to directly assemble raw uncorrected reads. Here we will therefore focus only on **correction-free assembly**. In this setting, assembly quality may become affected by e.g. chimeric reads and highly-erroneous regions^1^, as we will see next.

The `DASCRUBBER` program^2^ introduced the concept of read “scrubbing”, which consists of quickly removing problematic regions in reads without attempting to otherwise correct bases. The idea is that scrubbing reads is a more lightweight operation than correction, and is therefore suitable for high-performance and correction-free genome assemblers. `DASCRUBBER` performs all-against-all mapping of reads and constructs a pileup for each read. Mapping quality is then analyzed to determinate putatively high error rate regions, which are replaced by equivalent and higher-quality regions from other reads in the pileup. `MiniScrub` (LaPierre *et al.*, 2018) is another scrubbing tool that uses a modified version of `Minimap2` (Li, 2017) to record positions of the anchors used in overlap detection. For each read, `MiniScrub` converts anchors positions to an image. A convolutional neural network then detects and removes of low quality read regions.

Another problem that is even more upstream of read scrubbing is the computation of overlaps between reads. The storage of overlaps is disk-intensive and to the best of our knowledge, there has never been an attempt at optimizing its potentially high disk space.

In this paper we present two tools that together optimize the early steps of long-read assemblers. One is `yacrd` (Yet Another Chimeric Read Detector) for fast and effective scrubbing of reads, and the other is `fpa` (Filter Pairwise Alignment) which filters overlaps found between reads.

## 2 MATERIALS & METHODS

Similarly to `DASCRUBBER` and `MiniScrub`, `yacrd` is based on the assumption that low quality regions of reads are not well-supported by other reads. But unlike other tools, `yacrd` uses only approximate positional mapping information given by `Minimap2`, which avoids the time-expensive alignment step. This comes at the expense of not having base-level alignments, but this will turn out to be sufficient for performing scrubbing.

The `yacrd` algorithm proceeds as follows: all-against-all read mapping is performed using `Minimap2`, and a base coverage profile is computed for each read. Reads are split at any location where coverage drops below a certain threshold, and the low-coverage region is removed entirely. A read is completely discarded if a significant portion of its length (e.g. 40%) is below the coverage threshold. Figure 1 illustrates the process.`yacrd` time complexity is linear in the number of overlaps.

**Fig. 1.**
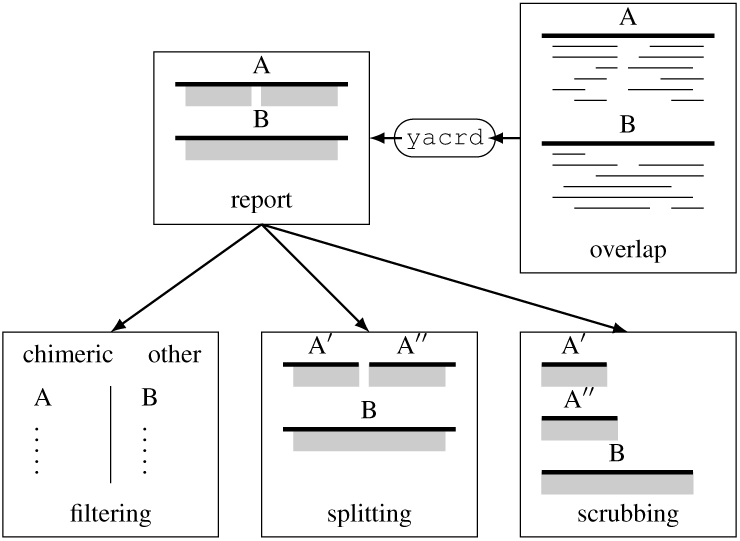
`yacrd` takes as input alignments and determines regions in reads that are sufficiently covered by alignments (grey boxes). `yacrd` then selectively perform i) chimeric reads filtering, ii) reads splitting, iii) keeping only high-confidence read regions (scrubbing).

`yacrd` performance is directly linked to the overlapper performance. We tuned a `Minimap2` parameter (the maximal distance between two minimizers, `-g` parameter) to find similar regions between reads and not to create bridges over low quality regions (see Supplementary Section 3). `yacrd` takes reads and their overlaps as inputs, and produces scrubbed reads, as well as a report.

`fpa` operates between the overlapper and the assembler. It is the first stand-alone tool capable of filtering out alignments based on a highly customizable set of parameters: e.g. alignments length, length of reads, reads names (see Figure 2). `fpa` can identify self-overlaps, end-to-end overlaps, containment overlaps, internal matches (when e.g. two reads share a repetitive region) as defined in (Li, 2016). `fpa` supports the PAF or BLASR m4 formats as inputs and outputs, with optional compression. `fpa` can also rename reads, generate an index of overlaps and output an overlap graph in GFA1 format.

**Fig. 2.**
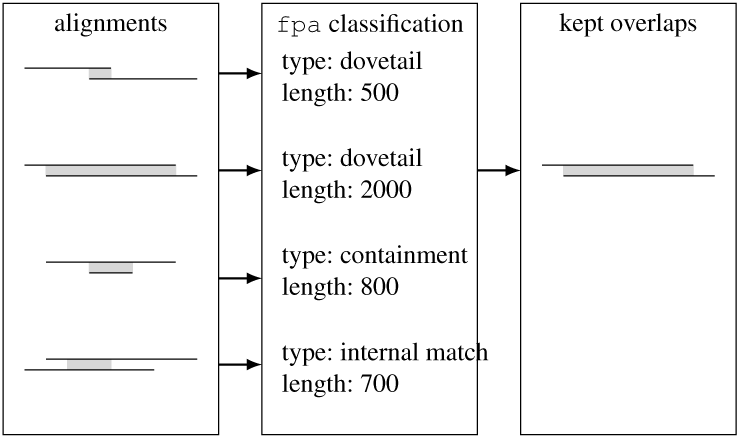
`fpa` analyzes each alignment separately and can filter baed on type (dovetail, containment, internal match), length, read length or read ID. In this example `fpa` kept only dovetail overlaps longer than 1 kbp.

`yacrd` and `fpa` are evaluated on 61 bacterial datasets from the NCTC database and from Maio *et al.* (2019) (details provided in Supplementary Section 1), representative of three long-read technologies: Oxford Nanopore (ONT), PacBio Sequel, PacBio RSII, as well as 3 eukaryotic datasets: *H. sapiens* chromosome 1 ONT ultra-long reads (from Jain *et al.* (2018)), *C. elegans* RSII reads^3^, and *D. melanogaster* Oxford Nanopore (ONT). All tools were run with recommended parameters (see Supplementary Section 2).

Scrubbed reads were then assembled using both `Miniasm` and `Wtdbg2` with recommended parameters for each sequencing technology. We compared `yacrd` with `DASCRUBBER`, and also executed `MiniScrub`. The latter took more than a day to run (without GPU support) on many bacterial datasets, and also it is only tailored to Nanopore data, therefore it was excluded from our benchmark. We used `Porechop`^4^ on all Nanopore datasets as a baseline number of adapters in reads.

## 3 RESULT & DISCUSSION

Table 1 presents the results of `yacrd` and `DASCRUBBER` on 3 representative bacterial datasets of each sequencing technology, as well as the 3 eukaryotic datasets.

**Table 1.**
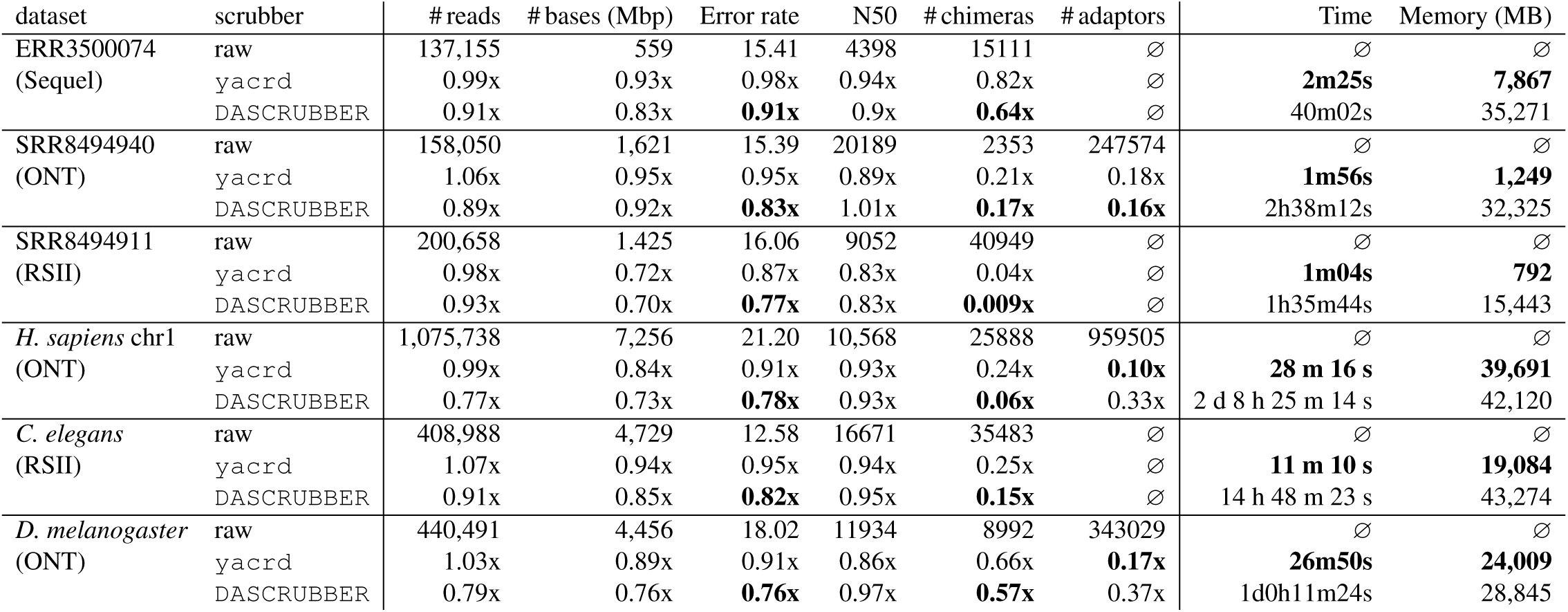
Performance of `yacrd` compared to `DASCRUBBER` on six representative datasets. # reads (resp. # bases) indicates the number of raw reads (resp. bases) or kept after scrubbing. Error rate is computed by running `samtools stats` after `bwa-mem` mapping of reads against reference. # chimera indicates the number of chimeric reads detected in the dataset using `Minimap2` (see Supplementary Section 4). # adaptors indicates the sum the number of adaptors found by `Porechop` at start/end of reads. Time and memory indicate respectively the wall-clock running time and peak RAM usage of `yacrd` and `DASCRUBBER`. Numbers in bold indicate best performance per dataset, nothing that deciding what is better for # reads, # bases and N50 is unclear.

| dataset | scrubber | #reads | #bases (Mbp) | Error rate | N50 | #chimeras | #adaptors | Time | Memory (MB) |
| --- | --- | --- | --- | --- | --- | --- | --- | --- | --- |
| ERR3500074<br>(Sequel) | raw | 137,155 | 559 | 15.41 | 4398 | 15111 | ∅ | ∅ | ∅ |
|  | yacrd | 0.99x | 0.93x | 0.98x | 0.94x | 0.82x | ∅ | **2m25s** | **7,867** |
|  | DASCRUBBER | 0.91x | 0.83x | **0.91x** | 0.9x | **0.64x** | ∅ | 40m02s | 35,271 |
| SRR8494940<br>(ONT) | raw | 158,050 | 1,621 | 15.39 | 20189 | 2353 | 247574 | ∅ | ∅ |
|  | yacrd | 1.06x | 0.95x | 0.95x | 0.89x | 0.21x | 0.18x | **1m56s** | **1,249** |
|  | DASCRUBBER | 0.89x | 0.92x | **0.83x** | 1.01x | **0.17x** | **0.16x** | 2h38m12s | 32,325 |
| SRR8494911<br>(RSII) | raw | 200,658 | 1,425 | 16.06 | 9052 | 40949 | ∅ | ∅ | ∅ |
|  | yacrd | 0.98x | 0.72x | 0.87x | 0.83x | 0.04x | ∅ | **1m04s** | **792** |
|  | DASCRUBBER | 0.93x | 0.70x | **0.77x** | 0.83x | **0.009x** | ∅ | 1h35m44s | 15,443 |
| *H. sapiens* chr1<br>(ONT) | raw | 1,075,738 | 7,256 | 21.20 | 10,568 | 25888 | 959505 | ∅ | ∅ |
|  | yacrd | 0.99x | 0.84x | 0.91x | 0.93x | 0.24x | **0.10x** | **28 m 16 s** | **39,691** |
|  | DASCRUBBER | 0.77x | 0.73x | **0.78x** | 0.93x | **0.06x** | 0.33x | 2 d 8 h 25 m 14 s | 42,120 |
| *C. elegans*<br>(RSII) | raw | 408,988 | 4,729 | 12.58 | 16671 | 35483 | ∅ | ∅ | ∅ |
|  | yacrd | 1.07x | 0.94x | 0.95x | 0.94x | 0.25x | ∅ | **11 m 10 s** | **19,084** |
|  | DASCRUBBER | 0.91x | 0.85x | **0.82x** | 0.95x | **0.15x** | ∅ | 14 h 48 m 23 s | 43,274 |
| *D. melanogaster*<br>(ONT) | raw | 440,491 | 4,456 | 18.02 | 11934 | 8992 | 343029 | ∅ | ∅ |
|  | yacrd | 1.03x | 0.89x | 0.91x | 0.86x | 0.66x | **0.17x** | **26m50s** | **24,009** |
|  | DASCRUBBER | 0.79x | 0.76x | **0.76x** | 0.97x | **0.57x** | 0.37x | 1d0h11m24s | 28,845 |

The main feature of `yacrd` is its short execution time, two orders of magnitude smaller than `DASCRUBBER`. In our tests, up to half of the execution time of `yacrd` is spent running `Minimap2`. Both scrubbers significantly reduce the number of chimeras in reads, with variable performance across datasets. `DASCRUBBER` tends to do a better job than `yacrd` at removing chimeras, but at the expense of discarding more reads and bases in reads. Across all datasets, the number of adapters is reduced by 72-94% with `yacrd` and 62-95% with `DASCRUBBER`. Read error rate is also marginally improved by 9-24%, yet this is not the main goal of these tools.

We next evaluate whether running `yacrd` results in higher-quality assemblies (Table 2). Both `yacrd` and `DASCRUBBER` overall improve NGA50 and reduce misassemblies (measured by the cumulative lengths of relocations) in `Miniasm` and `Wtdbg2` assemblies, compared to direct assembly of unscrubbed raw reads. Across nearly all datasets `yacrd` improves the NGA50 metric moreso than `DASCRUBBER` (except with the *C. elegans* dataset). We note that `Wtdbg2` contains steps that have a similar effect as `yacrd`, which explains why assembly metrics are not improved as significantly as with `Miniasm`.

**Table 2.** Performance of `yacrd` compared to `DASCRUBBER` on 64 datasets. For each dataset we report the ratio of metric X on the assembly done after yacrd (resp. `DASCRUBBER`) over metric X on the assembly done on raw reads, where X is either NGA50, largest alignment, or cumulative relocation length. For the bacterial datasets, we report the number of times the ratio is above 1, i.e. the number of times the metric on the assembly done after `yacrd` (resp. `DASCRUBBER`) is strictly larger (for NGA50 and longest alignment, and smaller for cumulative relocations length) than the metric on the assembly done on raw reads. For one of Sequel bacterial datasets, `DASCRUBBER` could not be executed.

| Bacterial datasets |  |  | NGA50 (# improved) |  | Longest Alignment (# improved) |  | Cumulative relocations lengths (# improved) |  |
| --- | --- | --- | --- | --- | --- | --- | --- | --- |
| Technology | # datasets | Scrubber | Miniasm | Wtdbg2 | Miniasm | Wtdbg2 | Miniasm | Wtdbg2 |
| Sequel | 22 | yacrd | 15 | 8 | 14 | 9 | 13 | 11 |
| Sequel | 21 | DASCRUBBER | 18 | 10 | 16 | 12 | 11 | 15 |
| ONT | 19 | yacrd | 18 | 11 | 15 | 12 | 11 | 9 |
| ONT | 19 | DASCRUBBER | 19 | 17 | 18 | 13 | 13 | 11 |
| RSII | 20 | yacrd | 19 | 13 | 17 | 16 | 14 | 16 |
| RSII | 20 | DASCRUBBER | 20 | 19 | 20 | 17 | 11 | 19 |
| Eukaryotic datasets |  |  | NGA50 (ratio/raw) |  | Longest Alignment (ratio/raw) |  | Cumulative relocations lengths (ratio/raw) |  |
| *H. sapiens* chr1 (ONT) | raw |  | 96 kbp | 1,643 kbp | 857 kbp | 9,267 kbp | 4,779 kbp | 4,088 kbp |
|  | yacrd |  | **5.14x** | **4.62x** | **4.65x** | **4.47x** | 1.11x | 0.94x |
|  | DASCRUBBER |  | 0.99x | 0.37x | 0.89x | 0.48x | **0.71x** | **0.6x** |
| *C. elegans* (RSII) | raw |  | 409 kbp | 537 kbp | 1,382 kbp | 1,382 kbp | 354 kbp | 245 kbp |
|  | yacrd |  | 1.04x | 1.04x | 1.34x | 1.23x | 0.8x | 0.8x |
|  | DASCRUBBER |  | **1.24x** | 1.04x | **1.37x** | **1.27x** | **0.55x** | **0.6x** |
| *D. melanogaster* (ONT) | raw |  | 838 kbp | 1,106 kbp | 5,193 kbp | 5,454 kbp | 664 kbp | 574 kbp |
|  | yacrd |  | **1.17x** | **0.93x** | **0.79x** | **1.03x** | 0.82x | 0.97x |
|  | DASCRUBBER |  | 0.75x | 0.62x | 0.65x | 0.62x | **0.83x** | **0.9x** |

On the *H. sapiens* and *C. elegans* datasets, `DASCRUBBER` reduces the total relocation length by a factor of 30-36% more than `yacrd`. However, given that all assemblies in Table 1 completed in less than an hour and `DASCRUBBER` took up to 2 days, running this tool on larger datasets would become a significant performance bottleneck. In Supplementary Section 3 we examine the behavior of `yacrd` across its parameter space. We observe that different parameters worked best for different technologies. Our recommended parameters are: `-g 500 -c 4` for ONT, `-g 800 -c 4` for PacBio RSII, `-g 5000 -c 3` for Sequel, where `-g` is the maximal distance between `Minimap2` seeds, and `-c` is the minimal coverage threshold for keeping portions of reads.

`fpa` reduced the size of the reads self-alignments file (PAF file produced by `Minimap2`) by 40-79% on the evaluated datasets, without any significant effect on quality assembly. As a consequence this reduces the memory usage of `Miniasm` by 13-67%. Other performance metrics are presented in Supplementary Table 1.

Finally, we examine the effect of **combining both** `yacrd` and `fpa`. We propose a pipeline based on `Miniasm` (see Supplementary Section 7) and show, across 5 datasets, that it consistently improves assembly contiguity, yields comparable assembly size, reduces mismatches and indels, reduces misassemblies, at the cost of a *≈* 2x increase in running time.

## Supporting information

assembly_metrics

scrubbing_result_on_read

## Acknowledgements

This work was supported by Inria and the INCEPTION project (PIA/ANR-16-CONV-0005). The authors thank Maël Kerbiriou for algorithmic help.

## Footnotes

1 https://dazzlerblog.wordpress.com/2015/11/06/intrinsic-quality-values/

2 https://dazzlerblog.wordpress.com/2017/04/22/1344/

3 github.com/PacificBiosciences/DevNet/wiki/C.-elegans-data-set

4 https://github.com/rrwick/Porechop/

