## Supplementary File for "yacrd and fpa: upstream tools for long-read genome assembly"

### 1 DATASETS

You can found in Supplementary File *dataset_information.csv* for each dataset is type (Nanopore, Pacbio Sequel, Pacbio RSII), shortest, longest reads, mean and median of read length, number of reads, number of bases, and the N50.

Dataset ERR3500074 and ERR3500075 is a merge in dataset named ERR3500074, ERR2651535 and ERR2651537 is a merge in dataset named ERR2651535, ERR3253077 and ERR3253092 is a merge in dataset named ERR3253077, ERR3253090 and ERR3253105 is a merge in dataset named ERR3253090, ERR3253086 and ERR3253101 is a merge in dataset named ERR3253086. They come from the same sample.

### 2 REPEATABILITY INFORMATION

#### 2.1 DASCRUBBER

DASCRUBBER commit number 0e90524 was used, and a custom pipeline was built using DASCRUBBBER-wrapper[1] as an inspiration, as well as recommendations from the authors: https://github.com/thegenemyers/DASCRUBBER/issues/7 and https://github.com/thegenemyers/DASCRUBBER/issues/20. See below (section 2.12) for the URL of the custom pipeline.

- DALIGNER commit fd21879
- DAMASKER commit 8243232
- DAZZ_DB commit 0bd5e07

#### 2.2 MiniScrub

We use version of commit 3d11d3e. We did not run MiniScrub in GPU mode so we followed te authors instructions for installation and run https://bitbucket.org/berkeleylab/jgi-miniscrub/.

#### 2.3 yacrd

We use version 0.5.1.

#### 2.4 fpa

We use version 0.5.

#### 2.5 bwa-mem

We use version 0.7.17-r1188.

#### 2.6 Minimap2

We use version 2.16-r922.

#### 2.7 Miniasm

We use version 0.3-r179

#### 2.8 Wtdbg2

We use version 2.3.

#### 2.9 Quast

We use version v5.0.2.

#### 2.10 Porechop

We use version 0.2.3_seqan2.1.1

#### 2.11 ReferenceSeeker

We use version 1.4

#### 2.12 Script and reproduction of analysis

All information to repeat our analysis can be found at this address
https://github.com/natir/yacrd-and-fpa-upstream-tools-for-lr-genome-assembly

[1] https://github.com/rrwick/DASCRUBBER-wrapper

### 3 YACRD PER-DATA-TYPE DETERMINATION OF PARAMETERS

yacrd is very dependent on the mechanism used to find common regions between reads. We rely on Minimap2 for this task. Minimap2 is based on short sequence seeds to find common regions between reads. In all-against-all alignment, it takes as parameter a distance between two seeds (-g, default: 10,000 bases). In yacrd we assume that regions with low seed coverage have low quality, and therefore need to be scrubbed. Yet with the default seed distance, it may happen that Minimap2 finds two consecutive seeds that correspond to two "good" read regions separated by one "bad" read region. Therefore this parameter needs to be tuned.

Another important parameter is the read coverage threshold to consider that a read region is of sufficient quality (yacrd parameter -c).

We have changed these two parameters as follows: i) the maximum distance between the two seeds from 50 to 2450 with a step of 100, ii) the minimum coverage before eliminating the region from 1 to 15 with a step of 1.

We evaluated the influence of these parameters on several metrics:

- Number of chimeric reads
- Number of reads
- Number of bases
- And in Miniasm and Wtdbg2 assemblies,
  - NGA50
  - Total length
  - Number of contigs
  - Number of indels per 100 kpb
  - Number of mismatches per 100 kbp

We ran this evaluation on *H. sapiens*, *C. elegans* and *E. coli* Pacbio dataset. The raw data is available in:

- *H. sapiens* (ONT ultra-long R9.4): https://gitlab.inria.fr/pmarijon/yacrd-and-fpa-upstream-tools-for-lr-genome- raw/master/data/yacrd_parameter_test_h_sapiens_ont.csv
- *C. elegans* (Pacbio P6-C4): https://gitlab.inria.fr/pmarijon/yacrd-and-fpa-upstream-tools-for-lr-genome-asse blob/master/data/yacrd_parameter_test_c_elegans_pb.csv
- *E. coli* (Pacbio Sequel): https://gitlab.inria.fr/pmarijon/yacrd-and-fpa-upstream-tools-for-lr-genome-assemb blob/master/data/yacrd_parameter_test_e_coli_pb.csv

For *H. sapiens* Nanopore dataset we find that a value of 500 for the -g parameter and 4 for the -c parameter optimizes the number of contigs in Miniasm assembly and NGA50, and remains reasonable across the other metrics. We therefore recommend to use this value for Nanopore data and we used it in all of our results.

For *C. elegans* PacBio dataset P6-C4, using a similar reasoning, optimal values are different and are 800 for the -g parameter and 4 for the -c parameter.

For *E. coli* PacBio Sequel dataset, using similar reasoning, optimal values are different and are 5000 for the -g parameter and 3 -c parameter.

We therefore used the above values for all datasets obtained with the same sequencing technology.

### 4 MAPPING OF SCRUBBED READS

To compute assembly quality metrics, we chose to evaluate each assembly against a well-chosen reference. For the eukaryota datasets and two bacteria dataset we use the following reference genomes: datasets SRR8494940 and SRR8494911 reference genome AE014075.1, *H. sapiens* chromosome 1 from GCA_000001405.27, for *D. melanogaster* we downloaded all chromosomes from this url [2] for *C. elegans* we download chromosomes from this url [3] . For the other bacterial datasets, we used the closest available reference found by `ReferenceSeeker`, using the `Wtdbg2` raw reads assembly as input to `ReferenceSeeker`. We discarded datasets for which the assembly size is twice larger (or twice lower) than the size of the chosen reference genome, or if the `ReferenceSeeker` ANI score is lower than 0.95.

For each dataset we mapped both scrubbed and raw reads against their a reference genomes with `bwa-mem` (we used `ont2d` preset for Nanopore reads, and `pacbio` preset for Pacbio reads). The mapping results were analyzed using a custom Python script[4] which reports the number of mapped reads, the sum of edit distances between each read and the matching reference sequence, the sum of positions of the genome mapped by a read, and the error rate.

To count the number of chimeric reads for each dataset, we remapped reads against each reference genome with `Minimap2` (we used `map-ont` preset for Nanopore reads, and `map-pb` preset for Pacbio reads). We analyzed the PAF (Pairwise Alignment Format) file outputted by `Minimap2` with a custom Python script[5]. This script parses a PAF file and associates to each read a list of pairs of starting/ending mapping positions. For each read, if two pairs of positions overlap in the corresponding list, they are merged. If, after merging, there remains more than one pair of positions, the read is marked as chimeric. To manage circular genomes we ignore reads with mapping positions near to the beginning/ending of the genome (within a distance of $\texttt{reference length} - 0.1 \times \texttt{reference length}$ from the beginning/ending).

To count the number of adapters in Nanopore reads we use `Porechop` with out any specific parameter and we sum the number of adapters at start and end of reads, we ignore the count of middle adapters.

Supplementary File *scrubbing_result_on_read.csv* value for all dataset, Pacbio dataset haven’t count of of adapters.

[2] `https://www.ebi.ac.uk/ena/data/view/AE014298.5,AE014134.6,AE013599.5,AE014296.5,AE014297.3,AE014135.4,CP007106.1,KJ947872.2&display=fasta`

[3] `https://www.ebi.ac.uk/ena/data/view/BX284601.5,BX284602.5,BX284603.4,BX284604.4,BX284605.5,BX284606.5&display=fasta`

[4] `https://gitlab.inria.fr/pmarijon/optimizing-early-steps-of-lr-assembly/blob/master/script/get_mapping_info.py`

[5] `https://gitlab.inria.fr/pmarijon/optimizing-early-steps-of-lr-assembly/blob/master/script/found_chimera.py`

### 5 QUALITY OF ASSEMBLY

To assess the quality of assemblies with and without scrubbing, we ran both `Miniasm` and `Wtdbg2` from scrubbed reads and raw reads with recommended parameters for each sequencing technology. After assembly we ran `Quast` with parameter `--min-identity 80.0`.

Supplementary File *assembly_metrics.csv* contains output metrics extracted from `Quast` reports and `ReferenceSeeker`,assembly for **all datasets, including the 61 bacterial datasets**. This is the raw data used to generate Table 2 in the main text.

In this file, we have reported the following assembly metrics:

- NGA50
- NGA50 with *extensive-mis-size* `Quast` parameter set to 10 kb
- Largest contig
- Largest alignment
- Total length
- Reference length
- Ratio of assembly length and reference length
- Indels per 100kb
- Mismatches per 100kb
- number of relocations
- number of translocations
- number of inversions
- cumulative lenght of relocations
- `ReferenceSeeker` ANI
- coverage

### 6 `FPA`

To evaluate `fpa`, we ran two different pipelines. The first one uses directly `Miniasm` without `fpa` and with recommended parameters. The second one runs `fpa` to filter out reads (`Minimap2` output is piped to `fpa` directly) before running `Miniasm` on filtered reads with recommended parameters. Using `fpa` we removed `internal match` and overlap shorter than 2000 (options `drop -i -l 2000`). This sort of overlap is ignored by `Miniasm` during the assembly step but is used during the read filtering step.

Table 1 shows the impact of using `fpa` on time, memory and assembly metrics. Using `fpa` decreases both disk usage and total computation time of downstream analysis while having no impact or a positive one on assembly metrics. Usage of `fpa` does not radically affect mapping wall-clock time and memory usage, but it reduces by 13% to 67% the memory usage and CPU time of the assembly step (the computation time of `fpa` is included in the mapping time). Moreover the size of the PAF file produced by `Minimap2` is reduced by 40% to 79 %.

| | Dataset | *C. elegans* | | *D. melanogaster* | | *H. sapiens* chr 1 | | SRR8494940 (ONT) | | SRR8494911 (RSII) | |
|---|---|---|---|---|---|---|---|---|---|---|---|
| | Pipeline | w/o `fpa` | `fpa` | w/o `fpa` | `fpa` | w/o `fpa` | `fpa` | w/o `fpa` | `fpa` | w/o `fpa` | `fpa` |
| Time (s) | Mapping | 3296 | 3247 | 3510 | 3659 | 1570 | 1558 | 26 | 29 | 23 | 24 |
| | Assembly | 297 | 139 | 782 | 186 | 103 | 50 | 4 | 2 | 2 | 1 |
| | Total | 3593 | 3386 | 4292 | 3845 | 1673 | 1608 | 30 | 31 | 25 | 25 |
| Memory | Mapping (GB) | 51 | 51 | 53 | 54 | 41 | 40 | 3 | 3 | 4 | 4 |
| | Assembly (Mbp) | 4788 | 2594 | 13836 | 5335 | 1797 | 587 | 52 | 45 | 33 | 22 |
| | **PAF size** | 32G | 9.5G | 54G | 11G | 8.9G | 3.2G | 141M | 82M | 85M | 38M |
| Assembly | # contigs | 168 | 150 | 423 | 381 | 184 | 216 | 5 | 5 | 8 | 9 |
| | NGA50 | 407821 | 438055 | 423007 | 455307 | 96225 | 106259 | 1450762 | 1246808 | 562741 | 292111 |
| | cum len of relocation | 291 mbp | 237 mbp | 4,779 mbp | 5,215 mbp | 664 mbp | 480 mbp | 1.6 mbp | 8.7 mbp | 4.4 kbp | 4.4 kbp |
| | length | 112248122 | 111641079 | 138733599 | 136623341 | 202082384 | 198386315 | 5147604 | 5283927 | 5394119 | 5395896 |
| per 100kb | # mismatches | 1893.44 | 1854.95 | 4233.35 | 4190.43 | 4089.56 | 4065.95 | 4341.81 | 4425.24 | 1862.72 | 1841.66 |
| | # indels | 7700.42 | 7628.39 | 5789.82 | 5742.05 | 6554.02 | 6534.92 | 5279.79 | 5376.03 | 7968.63 | 7945.11 |

**Table 1.** Impact of `fpa` on assembly using `Miniasm`. cum len of relocation correspond to cumulative length of relocation

### 7 COMBINATION OF YACRD AND FPA

To evaluate the effect of running both `yacrd` and `fpa`, we ran two different pipelines. The first one uses a standard `Miniasm` pipeline (called 'basic'): `Minimap2` plus `Miniasm` with recommended parameters. The second one (called 'extended') runs `yacrd` with best parameters for each dataset, then `Minimap2` with recommend parameter on scrubbed reads and pipes the results in `fpa` to filter out `internal matches` and overlaps shorter than 2000 (option `drop -i -l 2000`), and finally runs `Miniasm` on scrubbed reads with filtered overlap.

Table 2 shows how the integration of both `yacrd` and `fpa` in `Miniasm` pipeline ('extended' row) compares against standard `Miniasm` ('simple' row). Each pipeline is based on `Minimap2` so their memory usages are equivalent. The extended pipeline takes twice more time because `Minimap2` is run twice (once for `yacrd` and once for `Miniasm`). `Minimap2` is a time bottleneck in both pipelines.

The extended pipeline improves the quality of assemblies, in terms of NGA50, number of indels and mismatches per 100 kbp, and cumulative length of relocation (except for dataset SRR849411). It also decreases the number of contigs while keeping the total length of assemblies similar.

| Dataset | *C. elegans* | | *D. melanogaster* | | *H. sapiens* | | SRR8494940 (ONT) | | SRR8494911 (RSII) | |
|---|---|---|---|---|---|---|---|---|---|---|
| Pipeline | simple | extended | simple | extended | simple | extended | simple | extended | simple | extended |
| # contigs | 226 | 171 | 423 | 345 | 184 | 367 | 1 | 1 | 4 | 3 |
| NGA50 | 432112 | 451479 | 423007 | 715276 | 96225 | 488573 | 1450762 | 3775889 | 499610 | 1271550 |
| Largest contig | 5422030 | 4224860 | 8745435 | 5559611 | 15987693 | 6875897 | 5147604 | 5186180 | 1974889 | 4960107 |
| Largest alignment | 1231264 | 1527213 | 2396453 | 3053469 | 857015 | 4444801 | 1553466 | 3775889 | 1083557 | 1465922 |
| Total length | 114194187 | 110177189 | 138733599 | 134443509 | 202082384 | 202405973 | 5147604 | 5186180 | 5417095 | 5355278 |
| # indels per 100 kbp | 7842.91 | 7380.12 | 5789.82 | 5593.09 | 6554.02 | 6359.25 | 5279.79 | 5097.12 | 8011,42 | 7969,99 |
| # mismatches per 100 kbp | 1944.78 | 1720.16 | 4233.35 | 4052.42 | 4089.56 | 3884.23 | 4341.81 | 4113.01 | 1856,96 | 1844,42 |
| cum len of relocation | 354 mbp | 280 mbp | 4.779 mbp | 4.384 mbp | 664 mbp | 580 mbp | 384 kbp | 11 kbp | 4 kbp | 44 kbp |

**Table 2.** Impact of `yacrd` and `fpa` on assembly using `Miniasm`. Simple match to basic `Miniasm` pipeline and extend match to version with `yacrd` and `fpa`. cum len of relocation correspond to cumulative length of relocation.
